# A single nucleotide polymorphism genotyping panel for efficient genetic stock identification of the Teno river Atlantic salmon (*Salmo salar*) population complex

**DOI:** 10.1101/2024.10.04.616630

**Authors:** Helena Johansson, Jaakko Erkinaro, Morten Falkegård, Anders Foldvik, Panu Orell, Craig R Primmer, Victoria L Pritchard

**Author notes:** Corresponding author*: Craig Primmer, Viikinkaari 9, 00014, Helsinki, Finland.

## Abstract

A new single nucleotide polymorphism (SNP) panel for genetic stock identification in the Teno River Atlantic salmon fishery was developed, with a view to improving on an existing microsatellite panel. Twenty-two genetically differentiated reporting units were proposed based on population genetic analyses of 1212 individuals collected at 37 locations in the river and genotyped for > 33,000 genome-wide SNPs. A small subset of these SNPs was selected for GSI using an iterative process that considered their diversity and differentiation across reporting units. A genotyping-by-sequencing assay was developed to simultaneously genotype 180 of these GSI SNPs plus a sexing marker. This new SNP panel showed comparative GSI power to the microsatellite panel, with anticipated improvements in terms of cost, speed, and robustness and transferability across laboratories and genotyping platforms. Mixed stock analysis of the 2018 Teno River salmon catch using the new panel inferred all 22 reporting units to contribute to the fishery. Estimated catch proportions positively scaled with an independent estimate of reporting unit productivity: target spawning female biomass. This demonstrates the usefulness and efficiency of the 180 SNP panel for Atlantic salmon GSI in the Teno River system. If conducted on a regular basis, GSI can enable fine-tuning of management strategies to promote sustainable fishing.

## Introduction

Fisheries often consist of a mix of stocks – that is, they harvest individuals that originate from multiple genetically distinct populations. Key for effective management of such mixed-stock fisheries is accurate knowledge of how different source populations contribute to the catch (Begg *et al*., 1999; Palsbøll *et al*., 2007). This enables stock-specific management strategies aimed at safeguarding vulnerable populations while allowing continued harvest of healthier stocks (Crozier *et al*., 2004). Genetic stock identification (GSI) has been a mainstay of mixed-stock fisheries management for several decades (Anderson *et al*., 2008; Koljonen & McKinnell, 1996; Winans *et al*., 2004). The GSI approach involves first generating a genetic baseline of allele frequencies from populations that may contribute to the fishery. Samples from the mixed stock fishery are then genotyped with the same marker panel, and the likelihood of the baseline population allele frequencies producing the genotype in question in the fishery sample are estimated using individual assignment-based approaches or else overall catch proportions are estimated using mixture modelling (e.g. Anderson *et al*., 2008; Koljonen *et al*., 2005).

In recent years, single nucleotide polymorphisms (SNPs) have replaced microsatellites as the marker of choice in a range of population and conservation genetic applications, including GSI e.g. (Larson *et al*., 2014). Compared to microsatellites, which have commonly been typed by fragment size analysis, SNPs have advantages in terms of contemporary ease, speed and cost of genotyping, improved repeatability among laboratories and platforms, and a simpler mutation model allowing more straightforward interpretation of evolutionary and population genetic processes. One key benefit of SNPs compared to microsatellites for GSI is the potential availability of many thousands of easily genotyped loci, from which a sub-set of highly informative markers can be selected for a specific assignment task. This may enable more efficient stock assignment, improved resolution of more challenging population substructures (Larson *et al*., 2014) and/or the inclusion of SNPs to address different research and management questions (Aykanat *et al*., 2016; May *et al*., 2020).

Atlantic salmon, *Salmo salar* L., is an anadromous salmonid fish natively distributed across the north Atlantic coast of Europe and North America. Atlantic salmon are fished and managed across their range, and many wild populations are in long-term decline from a multitude of threats, some of which are poorly understood (Parrish *et al*., 1998). Management of the species can be politically and logistically complicated, spanning national borders and involving conflicts about rights of use (Crozier *et al*., 2004). One of the largest and most diverse wild Atlantic salmon populations in the world (Erkinaro *et al*., 2019) breeds in the large Teno (Norwegian: Tana, Sami: Deatnu) river system on the border of Finland and Norway in northernmost Europe. Previous genetic studies have revealed a highly structured population complex consisting of up to 28 genetically distinct, and temporally stable populations (Vähä *et al*., 2011, 2017, 2007, 2008), with evidence for fine-scale local adaptation (Mobley *et al*., 2019; Pritchard *et al*., 2018). Annual riverine catches have typically varied between 100-200 metric tons, but weak sea survival and returns in recent years suggest a sharp decline in most populations (Anon. 2024). To aid management, a microsatellite baseline (33 loci) was developed (Vähä *et al*., 2017) that included over 3,300 individuals from 36 locations within the river system. Based on these data, a maximum of 32 reporting units for GSI was proposed, with an estimated assignment accuracy of 36-100% depending on the reporting unit and analytical approach applied. The current salmon fishery management in the Teno River system is based on these genetic units and stock-specific status assessment of many of them (Anon. 2024).

The aim of this study was to develop a SNP genotyping panel for Teno River salmon GSI, with a view to improving on the microsatellite panel and producing a tool more robust to changes in genotyping technology, with a view to further improving and future-proofing the stock assessment system. We first genotyped original (Vähä *et al*., 2017) baseline samples, augmented with a set of more recent samples, for over 38,000 genome-wide SNPs. We used this SNP dataset to re-determine suitable reporting units and develop a 180 SNP genotyping-by-sequencing (GT-seq) panel for GSI. We compared the assignment efficacy of the new GT-seq panel with that of 10,000 genome-wide SNPs, and with the previous 33 microsatellite panel. The new SNP panel was then used to perform mixed stock assignment of salmon caught in the Teno River in 2018. As additional validation, GSI results were compared with an independent proxy of expected population size, target female biomass.

## Materials and Methods

### Baseline samples and datasets

Archived DNA samples from juvenile salmon in the (Vähä *et al*., 2017) baseline collection were used as the basis of the new SNP baseline (Figure 1, Table S1). Samples were quantified using a NanoDrop spectrophotometer (Thermo Scientific) and assessed for degradation using gel electrophoresis. For two of the 32 baseline reporting units in (Vähä *et al*., 2017) (Bavttájohka and Anarjohka) there were either insufficient samples, the pre-extracted DNA was completely degraded or the DNA did not pass the minimum quality requirements for SNP genotyping. To augment the baseline, DNA was extracted and quantified for juvenile samples collected in 2014 from four localities, Tsarsjoki, Lakšjohka, and the Lakšjohka tributaries Deavkkehanjohka and Gurtejohka. Good quality DNA samples with a minimum concentration of 20ng/ul were sent to the Center for Integrative Genetics (CIGENE), Ås, Norway, for SNP typing on a custom 60,250 (60K) Atlantic salmon SNP genotyping array. All subsequent merging and quality control of SNP datasets was performed using Plink 1.90 (Chang *et al*., 2015).

**Figure 1.**
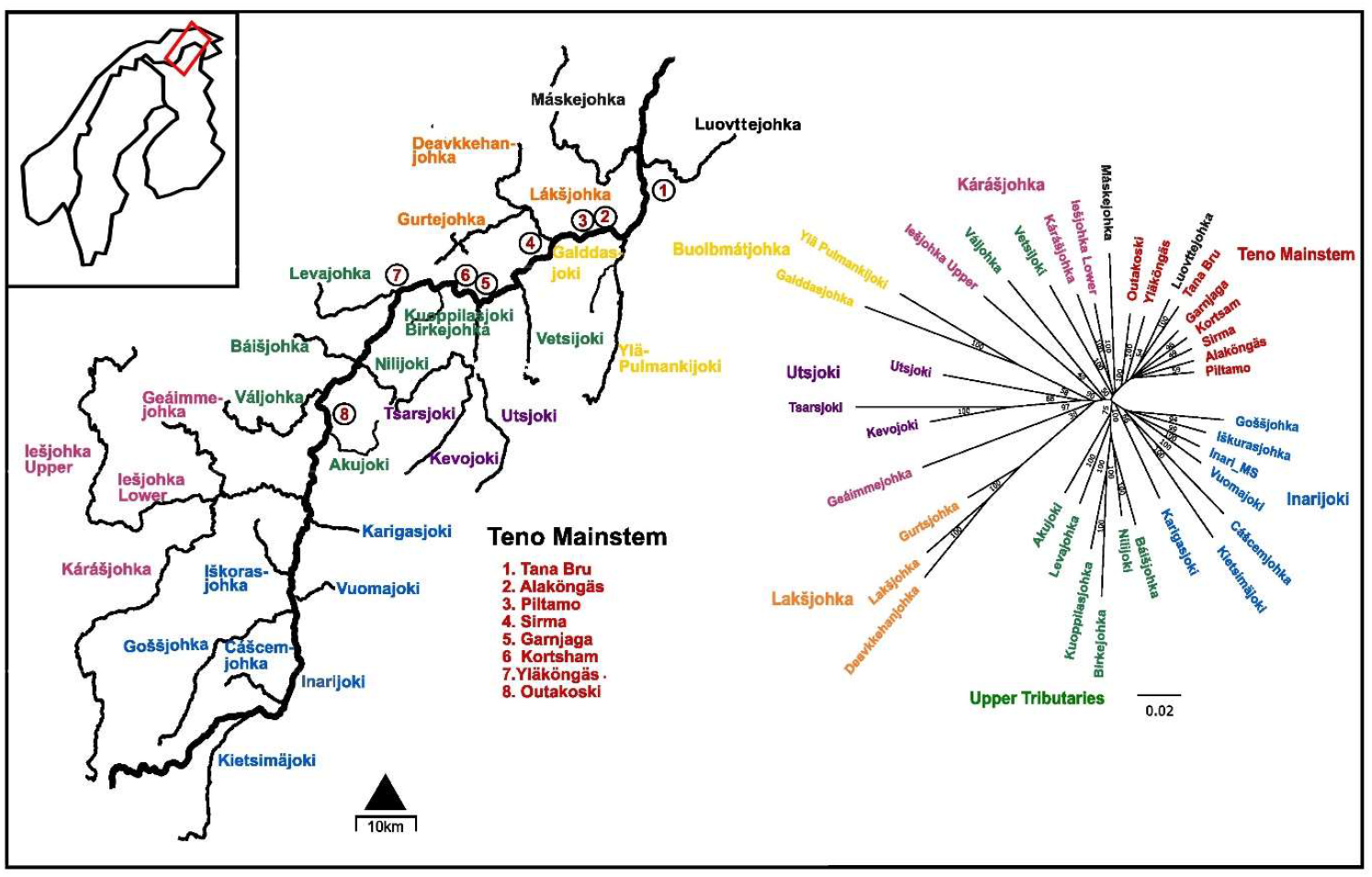
Map of the Teno river system baseline sampling localities and Neighbour-joining tree using Prevosti’s distance on 17,546 SNPs. Geographic regions indicated by colour in map and in tree.

To further extend the data set, we added genetic data generated for previous studies of Teno River salmon (Barson *et al*., 2015, Pritchard *et al*., 2018): adult genotypes from one locality missing from the 60K juvenile baseline (Máskejohka) and additional juvenile genotypes – also part of the original baseline -from ten localities. These samples had been screened using a 220K SNP array, which shares 38,669 named SNPs with the 60K array. Forty-one individuals from Pritchard *et al*. (2018) were typed on both arrays, enabling genotyping repeatability be examined. After merging the 60K and 220K datasets across the 38,669 markers, we removed 364 SNPs with which had > 2/41 mismatching genotypes across arrays and four SNPs with known off-target variants. Subsequently we applied the following filters: SNP minor allele frequency >0.05; SNP missing genotype frequency < 0.03; individual missing genotype frequency < 0.05. Finally, putative full-sibs were identified and removed based on genome-wide patterns of identity-by-descent (--genome) within and across collection locations.

### Baseline population substructure

Before analyses of population structure, the baseline SNP dataset was further filtered to prune neighbouring SNPs in high linkage disequilibrium with one another (--indep 50 5 1.5). Population genetic analyses were carried out either in the R statistical environment (R Core Team, 2014), or in the Linux shell.

Overall and pairwise F_ST_s among samples from different locations were estimated using *StAMPP* (Archer *et al*., 2017). Significance of pairwise F_ST_ was assessed using 20,000 bootstrap replicates across loci and correcting for multiple testing using the Bonferroni procedure with alpha = 0.05. A neighbour-joining (NJ) tree based on Prevosti’s distance (Prevosti *et al*., 1975), was created and bootstrapped 100 times using *poppr* (Kamvar *et al*., 2014) and the tree plotted using the package *ape* (Paradis & Schliep, 2019). Prevosti’s distance is a model free distance suitable for use with SNP data with missing genotypes.

*Admixture* (Alexander *et al*., 2009) was used to identify the most likely number of genetically distinct ancestral populations present in the LD pruned baseline SNP dataset, independent of sampling location. Five replicates of the Admixture analysis were run, each starting from a different random number seed, allowing between 1 and 30 genetically distinct populations in the dataset. The most likely number of populations was determined using the software’s cross-validation procedure. Finally, genetic variation within the dataset was also explored using principal component analysis followed by graphical visualization. PCA was performed using the package LEA in R (Frichot & Francois 2015), with number of retained PCs based on Tracy-Widom test results. The combined results from the Admixture and PCA analyses, interpreted with reference to (Vähä *et al*., 2017) were used to select suitable new reporting units for GSI of the Teno River fishery.

### Developing and testing of the GT-seq SNP panel

To identify a sub-set of SNP markers for efficient mixed stock assignment, highly variable SNPs (MAF > 0.45), were first extracted from the full baseline dataset using PLINK 1.9. Following this, genotypes from eight highly differentiated reporting units (Kevojoki, Tsarsjoki, Utsjoki, Lakšjohka, Deavkkehanjohka, Gurtejohka, Yla-Pulmankijoki, Galddasjoki) were removed, as inclusion of these sites greatly skewed the *F*_ST_ distribution for SNPs, while assignment efficiency should be high for these reporting units regardless of the specific markers used. Closely linked SNPs in this new dataset were LD pruned (indep --50 5 1.5), and the remaining SNPs were ranked by calculating *F*_ST_ over the remaining reporting units. The 96 highest ranking SNPs were retained as the initial GSI panel. Filtering for highly variable SNPs retains a markers set that is expected to be informative across multiple reporting units, and subsequent selection of SNPs with the highest *F*_ST_ across units is expected to maximise accuracy for GSI (Ackerman *et al*., 2011; Storer *et al*., 2012). This initial SNP panel was tested for its assignment efficacy using the ‘leave-one-out’ individual re-assignment approach implemented in the R package *rubias* (Moran & Anderson, 2019). Markers were then added to improve discrimination among target subsets of reporting units by selecting additional SNPs with high MAF and high *F*_ST_ across those units alone. This process was performed iteratively, aiming for a final panel of 220-230 candidate GSI SNPs which was intended to translate into working panel of approximately 200 SNPs. As an additional check to minimize marker redundancy due to linkage, the positions of final candidate SNPs were inspected to exclude any markers within 10,000 bp of one another. Finally, SNPs linked to several genes identified as locally adaptive candidates in the Teno River (Pritchard *et al*., 2018) and the *sdY* sex marker (Yano *et al*., 2013) were added to the panel.

Primers for GT-Seq (Campbell *et al*., 2015) were designed to amplify a 120-155bp DNA sequence surrounding each of the target SNPs using BatchPrimer3 (Rozen & Skaletsky 1998;Untergasser *et al*., 2012), with the following parameters: Tm 58-62°C optimum 60°C; GC% 28-68; all other parameters default. Primers were not assessed *in silico* for dimerization potential.

All multiplex PCR was performed in 12ul reactions containing the following: 6ul Qiagen Multiplex PCR mix, 2ul DNA at a concentration of 10-50 ng/ul, volume of multiplexed diluted primers to yield the target concentration of each primer (see below); molecular grade water. Thermocycling conditions were: 98°C for 2 min, 20x [98°C 10s, 62°C 30s, 72°C 15s], 7C for 10 min. PCR products were tagged with individual-specific barcode indices, pooled, and sent to the DNA Sequencing and Genomics Laboratory, University of Helsinki for library preparation and sequencing. Products were single-end sequenced using Illumina150 cycle kits on an Illumina HiSeq or MiSeq. Sequences were demultiplexed by barcode index using Generate FASTQ (Illumina) and 3’ trimmed on the basis of indexing adaptor sequence using cutadapt 3.2 (Martin, 2011). Trimmed sequences were aligned using bwa mem (BWA 0.7.17, Li & Durbin, 2010) to the *S. salar* reference genome (ICSASG_v2) with an appended *S. salar* sdY sequence (Genbank Accession KT223110.1). Variants at targeted sites were called using samtools mpileup piped to bcftools call (samtools 1.12, Li & Barrett, 2011). The output VCF file was imported into Plink 1.9 with genotypes filtered for a minimum GQ of 15 and a minimum sequencing depth of 8. For genetic sexing, sdY sequencing depth was corrected for overall sequencing depth for each individual. Individuals were then assigned as male (sdY present), female (sdY absent) or unknown on the basis of thresholds chosen by examining the distribution of corrected sdY sequencing depths over all individuals.

Two rounds of multiplex PCR optimization were performed for the GT-Seq panel, each using 16 samples as the amplification target. For the first optimization round, all primers were combined in equimolar concentrations (0.03 µM) in a single reaction. The output fastq files for the 16 test fish were merged, and a custom bash script was used to count number of sequence reads for each possible forward-reverse primer combination. This information was used to identify primers producing dimer sequence, and primers were split into two multiplex pools that minimized within-pool dimer formation. The subsequent optimization round was used to check pool composition and to adjust primer concentrations (final range 0.03-0.06 µM) with the aim of roughly equalizing sequencing coverage across SNPs. After these two rounds of optimisation the SNP panel was used to genotype 74 baseline individuals to check for genotyping success and consistency of genotype calling between GT-Seq and the 60K SNP array.

### Assignment efficacy of SNP panel

The accuracy of GSI across identified reporting units was tested using two methods implemented in *rubias*: the ‘leave one out’ approach described above, and a ‘100% mixture’ approach. In the latter analysis, baseline allele frequencies are used to simulate the genotypes of fish deriving from different populations and those fish are then assigned back to the full baseline. We simulated 500 fish for each population and performed 100 replicates. To compare the performance of the SNP panel to that which could be achieved by a much denser sampling of genetic variation across the genome, we randomly subsampled 10,000 SNPs from the baseline dataset and examined assignment power of this dataset via a leave-one-out analysis implemented in gsi_sim, a Linux command line version of *rubias* suitable for large datasets (https://github.com/eriqande/gsi_sim, (Anderson, 2010; Anderson *et al*., 2008).

To assess the robustness of assignments with smaller numbers of SNPs from the SNP panel, and thus assess the effect of missing genotypes on GSI reliability, random subsets of the 180 SNPs (60, 80, 100, 120, 140 and 160 loci) were tested in ‘leave-one-out’ analyses against the final set of reporting units.

### Comparison with the microsatellite baseline

We compared the efficacy of the new SNP GSI panel with that of the previous microsatellite panel. To do this, we used the existing microsatellite baseline, which comprises 3,323 juvenile and adult salmon from 36 sites sampled within the Teno River system and typed at 33 microsatellite loci. Sampling and molecular protocols for this dataset can be found in (Vähä *et al*., 2011, 2017, 2007, 2008). We compared the assignment performance of the SNPs and microsatellites using the *rubias* leave-one-out procedure under two reporting unit scenarios: the 22 units proposed in this paper and the 32 reporting units (two not represented in the SNP baseline) used by Vähä *et al*., 2011.

### Mixed stock assignment and validation

In 2018, scale samples from salmon caught in the Teno River were collected by local fishers, coordinated by Natural Resources Institute Finland (LUKE, Finland) and the Norwegian Institute for Nature Research (NINA, Norway). Fishers also recorded standard data on the fish including length, mass, morphologically determined sex and visually assessed whether they were wild or aquaculture escapee origin (Erkinaro et al. 2010). We extracted DNA from 1-2 scales for each fish (excluding two putative escapees) using a high-salt method, quantified extractions using NanoDrop, and standardized to a concentration of 10-50ng/µl. Samples from which sufficient DNA could be obtained were genotyped for the 180 GSI SNPs and sex marker as described above. Individuals missing genotypes at >60 loci were removed from the data set. Finally, we identified duplicate samples on the basis >of 90% genotype match with another sample using the close_matching_samples function in *rubias* and in each case excluded the duplicate with the fewest missing genotypes.

To estimate the proportional contribution of each of the 22 reporting units to the 2018 Teno River catch, we used *rubias* to perform mixed stock analysis against the SNP baseline. We implemented the maximum likelihood approach with the default options of a uniform prior, 20000 sweeps and the first 1000 sweeps discarded as burn-in. We computed 95% credible intervals from the MCMC traces of mixing proportions.

To indirectly validate the results of the mixed stock analysis, we used location-specific figures for target spawning female biomass provided in (Falkegård *et al*., 2014). This is the biomass of females required to produce the target number of eggs for each location, which is estimated by modelling habitat suitability and egg density estimates and taking stock-specific fecundity into account. It can be considered a proxy for the expected numbers of breeding adults returning to different parts of the Teno River system. In (Falkegård *et al*., 2014), tributaries were grouped by geography into somewhat different groups than the reporting units that we use here. For the purpose of comparison, the mixed stock analysis proportions were combined to match the (Falkegård *et al*., 2014) geographic groupings, and female biomass (kg) estimates were transformed into relative proportions of the total for each group (Table S5). A Pearson correlation was performed to compare the estimated female biomass proportions to the proportions from the mixed stock assignment using cor.test in base R. Although not comparing the same thing, as one is based on individuals, and the other on biomass, it provides an indirect assessment of the realism of the GSI proportion estimates.

## Results

### Genetic baseline

The initial genetic baseline comprised 1,309 samples, collected between 2006 and 2014 from 37 locations in the Teno River (mean per location 35.3, range 11-67) and genotyped at 33,139 post-QC SNPs. Following removal of putative full siblings and LD pruning, a total of 1,212 individuals genotyped at 17,546 loci (Table S1) were used to investigate baseline population structure.

Global F_ST_ demonstrated substantial genetic structuring among sample sites across the Teno River system (F_ST_ =0.055, *P*=0.0099). Pairwise F_ST_s ranged from 0.0005 (between Piltamo and Alaköngäs) to 0.1953 (between Deavkkehanjohka and Galddasjoki) and all were statistically significant, with the exception of the comparison between Piltamo and Alaköngäs (Table S2). Nevertheless, the generally very low pairwise F_ST_ among samples from Tana Mainstem localities (Tana bru, Alaköngäs, Piltamo, Sirma, Garnjarga, Kortsham, Yläköngäs, Outakoski) indicated only weak genetic differentiation across most of the Teno River mainstem.

The neighbour-joining tree reflected the pairwise F_ST_ results, showing a strong geographic signal across sampling locations (Figure 2), with strong bootstrap support at many nodes.

**Figure 2:**
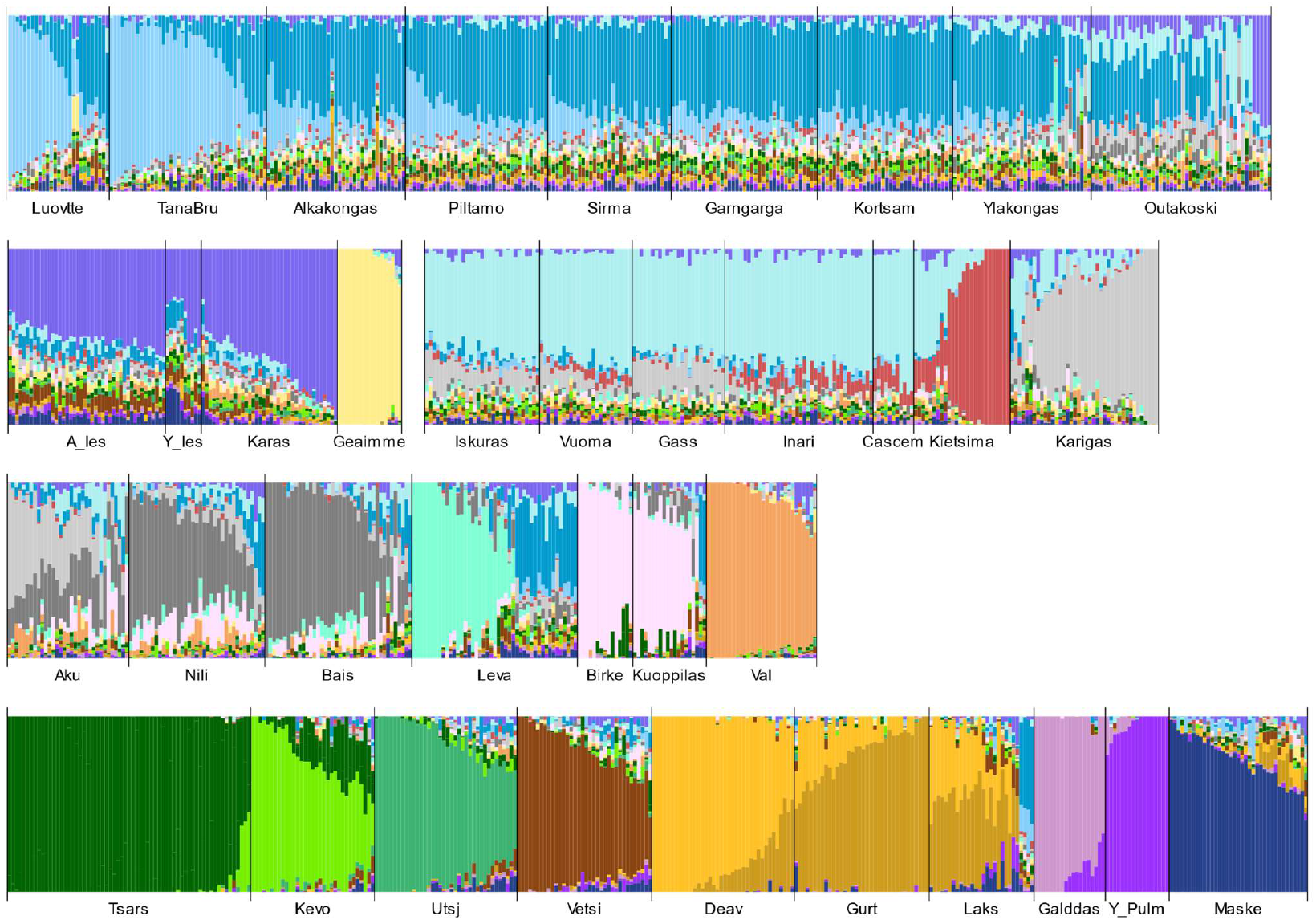
Results of Admixture analysis of Teno River Atlantic salmon individuals. Each column represents an individual fish, ordered by sampling location, indicated on the y-axis. Colours indicate inferred proportion of ancestry (Q) from each of 21 possible ancestral clusters.

Admixture inferred 21 ancestral clusters within the dataset (Figure 3), with a geographic signal in line with the F_ST_ analysis and the neighbour-joining tree and indicative of gene flow among neighbouring locations. The Teno River mainstem and its two large headwater systems (Inarijoki and Kárášjohka) were separately dominated by three large ancestral clusters. The uppermost sample from the mainstem (Outakoski) showed substantial mixing among these clusters. A fourth genetic cluster dominated the lowermost mainstem sample, Tana Bru and the nearby tributary Luovtejohka. Ten smaller tributaries (Karigasjoki, Máskejohka, Galddasjoki, Ylä-Pulmankijoki, Utsjoki, Kevojoki, Váljohka, Kietsimäjoki, Vetsijoki and Geáimmejohka) and two pairs of neighbouring tributaries (Kuoppilasjoki and its tributary Birkejohka and Báišjohka/Nilijoki) were dominated by their own unique ancestral clusters. Akujoki had a mixed ancestry largely from the neighbouring Karigasjoki and Báišjohka/Nilijoki clusters. Two ancestral clusters were inferred for samples from Lakšjohka and its two small tributaries Gurte- and Deavkkehanjohka; examination of the underlying data suggested that this was likely caused by a segregating chromosomal rearrangement not accounted for by LD pruning. The large sample from Tsarsjoki was split into two clusters, which corresponded to year of collection likely reflecting differences in their sampling location in the tributary: these were amalgamated for subsequent analyses.

**Figure 3.**
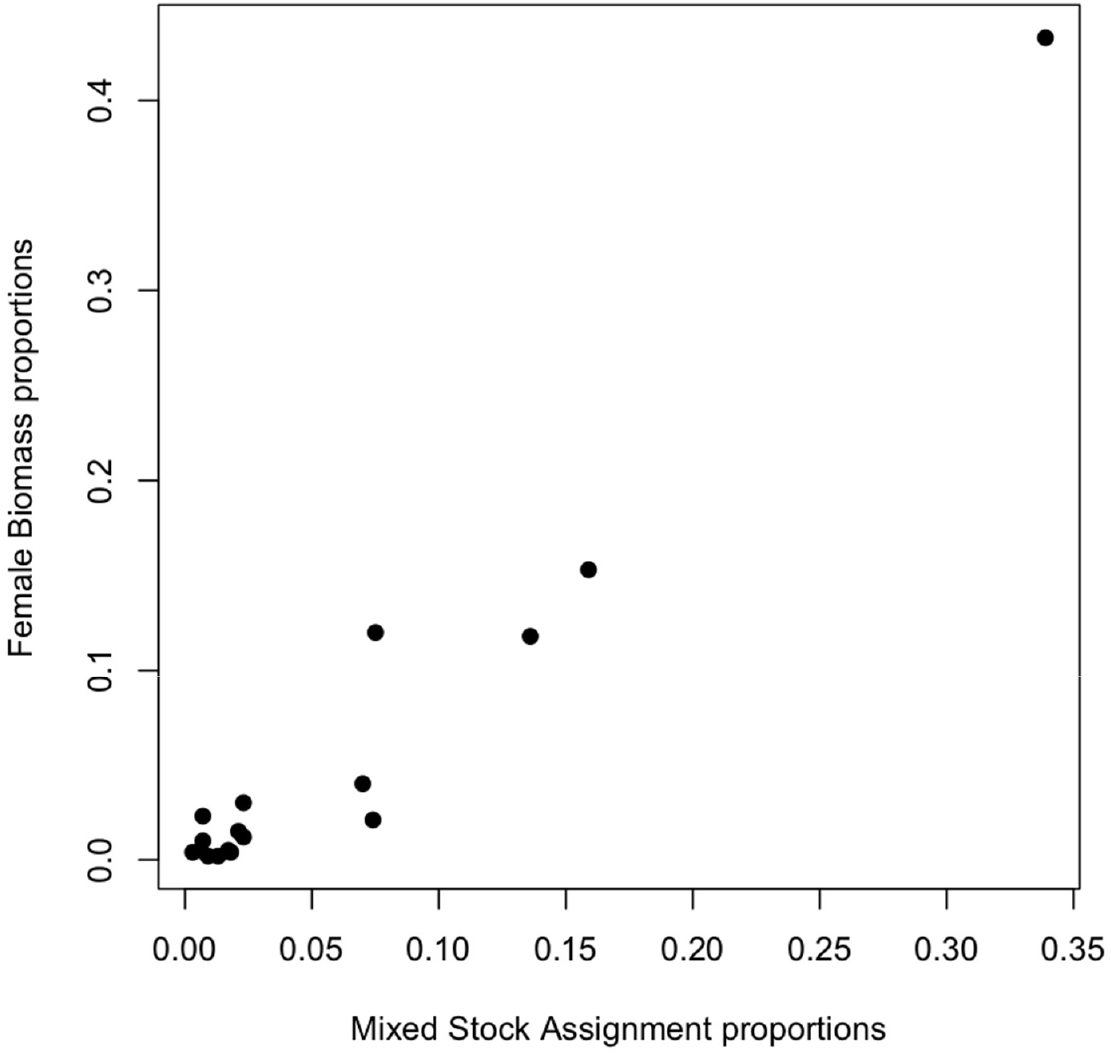
The proportion of female biomass per river/tributary (estimated spawning target converted to required female biomass from Falkegård et al. (2014)) plotted against the estimated catch proportion in the mixed fishery for the same reporting units.

Twenty three principal component were retained in the PCA. Graphical examination of the position of individuals along these PCs (not shown) supported the conclusions of the Admixture analysis with three exceptions: the Outakoski sample partly separated from the rest of the Teno River mainstem samples (Alaköngäs, Piltamo, Sirma, Garnjarga, Kortsham, Yläköngäs) along PC5, probably reflecting the genetic mixing in this location, the Iesjohka and Karasjohka samples were separated along PC23, and Akujoki was partly separated from other samples along PC21. Based on the combined Admixture and PCA results, together with consideration of Vähä *et al*., 2017, 22 reporting units were defined for GSI (Table 2), with Lakš/Gurte/Deavkkehanjohka considered a single reporting unit but Outakoski, Akujoki and Kárášjohka treated as separate reporting units.

In some tributaries, a subset of juveniles were assigned very high mainstem ancestry by the Admixture analysis and clustered with mainstem fish in the PCA. Examination of available capture locations showed that these 24 fish were sampled near tributary mouths; they were therefore considered to be migrants from the Teno River mainstem breeding population rather than part of the genetically distinct tributary breeding population and were removed from the final GSI baseline.

### SNP panel development

Primers were successfully designed for a final set of 223 candidate GSI SNPs, plus one marker each for candidate adaptively important genes *vgll3, six6, numa1* and *btg3* and genotypic sex (sdY). nFollowing initial optimization and tests, SNPs were excluded due to complete non-amplification (n=11), >10% mismatching genotypes between GT-Seq genotypes and SNP array genotypes (n=16, mostly missing heterozygotes), and >40% missing genotypes (n= 21). One pair of primers amplified two SNPs that were both included in the GSI panel. Primer sequences for the 180 SNPs in the final GSI panel plus the sdY locus are provided in the Supplementary Materials.

### Assignment efficacy

The leave-one-out analysis using the 180 SNP panel correctly assigned 86% of individuals to their reporting unit of origin. (Table 1). Thirteen out of the 22 reporting units had assignment accuracies between 90-100%, and only Outakoski, Tana Bru and Akujoki reporting units had assignment success < 70%. Mis-assigned fish were generally assigned to geographically adjacent reporting units (Table 1). Assessment of assignment efficacy using the 100% mixture approach gave very similar results, with slightly better accuracy inferred for all reporting units (93% of simulated individuals correctly assigned). (Table S2).

**Table 1.** Teno River Atlantic salmon GSI correct assignment rates and main unit of mis-assignment using the 180 SNP panel, and comparison of correct assignment achieved using microsatellites.

| Reporting unit | SNPs |  |  | Microsatellites |  |
| --- | --- | --- | --- | --- | --- |
|  | Baseline sample size | % correct assignment | Largest misassignment to: | Baseline sample size | % correct assignment |
| ***Teno Mainstem*** |  |  |  |  |  |
| TanaBru/Luovtejohka | 69 | 61% | Teno Mainstem (37%) | 127 | 66% |
| Middle Mainstem * | 220 | 83% | Tana Bru (12%) | 446 | 83% |
| Outakoski | 48 | 52% | Teno MS (18%); Inari HW (15%) | 91 | 54% |
| ***Lower Tributaries*** |  |  |  |  |  |
| Máskejohka | 37 | 86% | Teno Mainstem (8%) | 121 | 97% |
| Vetsijoki | 36 | 92% | Iešjohka (5%) | 212 | 96% |
| Ylä-Pulmankijoki | 17 | 100% | - | 279 | 99% |
| Galddasjoki | 19 | 100% | - | 85 | 99% |
| Utsjoki | 38 | 97% | Vetsijoki (3%) | 129 | 99% |
| Tsarsjoki | 65 | 97% | Kevojoki (3%) | 196 | 97% |
| Kevojoki | 33 | 100% | - | 165 | 99% |
| Lakš/Deavkke/Gurtsjohka | 98 | 97% | Karigas; Máske; Vetsi (1% each) | 71 | 100% |
| ***Upper Tributaries*** |  |  |  |  |  |
| Levajohka | 31 | 74% | Outakoski (6%); Báiš/Nilijoki (6%) | 51 | 69% |
| Báišjohka/Nilijoki | 73 | 85% | Akujoki (6%) | 106 | 77% |
| Akujoki | 33 | 70% | Báišjohka/Nilijoki (12%) | 53 | 40% |
| Karigasjoki | 40 | 83% | Akujoki (10%) | 75 | 79% |
| Kuoppilasjoki/Birkejohka | 33 | 91% | Báišjohka/Nilijoki (10%) | 112 | 91% |
| Váljohka | 30 | 93% | Inari HW (7%) | 270 | 94% |
| ***Kárášjohka/Iešjohka lower*** |  |  |  |  |  |
| Iešjohka | 54 | 93% | Kárášjohka (5%) | 201 | 89% |
| Kárášjohka | 40 | 83% | Iešjohka (3%); Vetsijoki (3%) | 270 | 94% |
| Geáimmejohka | 18 | 94% | Akujoki (5%) | 52 | 92% |
| ***Inari Headwaters*** |  |  |  |  |  |
| Inari Headwaters** | 132 | 91% | Karigasjoki (4%) | 177 | 85% |
| Kietsimäjoki | 26 | 73% | Inari HW (15%) | 61 | 74% |

\*Teno River Mainstem reporting group: Alaköngäs, Piltamo, Sirma, Garnjaga, Kortsham and Yläköngäs
\*\*Inari Headwaters reporting group: Inari Mainstem, Cášcemjohka, Vuomajoki, Goššjohka, Iškurasjohka

**Table 2.** Mixed stock analysis of the Teno River mainstem Atlantic salmon fishery catch in 2018.

| Reporting Unit | Estimated mixture proportion | Lower 95% CI | Upper 95% CI |
| --- | --- | --- | --- |
| ***Tana Mainstem*** |  |  |  |
| TanaBru/Luovtejohka | 0.029 | 0.0214 | 0.0378 |
| Teno Mainstem* | 0.251 | 0.2335 | 0.2689 |
| Outakoski | 0.059 | 0.0480 | 0.0700 |
| ***Lower Tributaries*** |  |  |  |
| Máskejohka | 0.023 | 0.0171 | 0.0297 |
| Vetsijoki | 0.074 | 0.0641 | 0.0851 |
| Váljohka | 0.021 | 0.0155 | 0.0265 |
| Ylä-Pulmankijoki | 0.007 | 0.0044 | 0.0109 |
| Galddasjoki | 0.001 | 0.0003 | 0.0030 |
| Utsjoki | 0.019 | 0.0144 | 0.0250 |
| Tsarsjoki | 0.019 | 0.0140 | 0.0243 |
| Kevojoki | 0.032 | 0.0253 | 0.0386 |
| Lakš/Deavkkehan/Gurtsjohka | 0.007 | 0.0044 | 0.0107 |
| Geáimmejohka | 0.013 | 0.0086 | 0.0171 |
| ***Upper Tributaries*** |  |  |  |
| Levajohka | 0.018 | 0.0133 | 0.0237 |
| Báišjohka/Nilijoki | 0.023 | 0.0171 | 0.0285 |
| Akujoki | 0.009 | 0.0048 | 0.0129 |
| Karigasjoki | 0.017 | 0.0120 | 0.0232 |
| Kuoppilasjoki/Birkejohka | 0.006 | 0.0031 | 0.0090 |
| ***Iešjohka Headwaters*** |  |  |  |
| Iešjohka Upper/Lower | 0.136 | 0.1227 | 0.1507 |
| Karášjohka | 0.075 | 0.0639 | 0.0854 |
| ***Inari Headwaters*** |  |  |  |
| Inari Headwaters** | 0.159 | 0.1445 | 0.1733 |
| Kietsimäjoki | 0.003 | 0.0009 | 0.0050 |

\*Teno Mainstem reporting group: Alaköngäs, Piltamo, Sirma, Garnjaga, Kortsham and Yläköngäs
\*\*Inarijoki reporting group: Inarijoki Mainstem, Čáčcemjohka, Vuomajoki, Goššjohka, Iškurasjohka

When testing both with the leave-one-out analysis, the 180 SNP panel performed almost as well as 10,000 genome-wide SNPs to assign samples to reporting unit, indicating that it is well captures genome-wide patterns of population substructure (Table S3). Overall assignment accuracies decreased gradually with each reduction in SNP number for sub-sets of the 180 GT-Seq panel however even with 100 SNPs 80% of fish were assigned back to their correct reporting unit (Table S3). Assignment success for the different size panels generally showed consistent patterns across the same reporting units.

The same two analyses applied to the larger microsatellite baseline split by the new reporting units demonstrated comparable assignment efficacy of the microsatellites (88% correct assignment using the leave-one-out approach, 90% correct assignment for the 100% mixture approach, Table 1 & S2). Similarly to the SNP panel the lowest assignment accuracy was for the Outakoski, Tana Bru and Akujoki reporting units.

### Mixed stock analysis

Of 2,116 scale samples received from the 2018 mixed stock fishery, 187 failed genotyping and a further 16 were identified as duplicate samples. This left 1,913 available for mixed stock analysis against the 22 reporting unit SNP baseline (mean proportion of missing genotypes = 0.04, range = 0 – 0.32). Three of the largest reporting groups of the mixed stock analysis (Table 2) were each assigned >10% of the fishery samples, totalling almost 55% of the total catch: Teno River mainstem (25.1%), Inarijoki (15.9%) and Iešjohka (13.6%). Three additional reporting groups were assigned >5% each: Kárášjohka (7.5%); Vetsijoki (7.4%) and Outakoski (5.9%) with the remaining 16 reporting groups each being assigned 3.2% or less of the catch (24.6% of the total catch). The MSA proportions were highly and significantly correlated with the proportions of female biomass target abundance (Figure 3, r= 0.97, p<0.000). About 9% fewer fish were assigned to Tana Bru/Tana Mainstem/Outakoski than expected based on the female biomass target abundance proportions, similarly, 4.5% less fish were assigned to Kárášjohka than expected. Conversely, about 5% more fish were caught from Vetsijoki, and 3% more from Utsjoki/Tsarsjoki/Kevojoki than expected from the target abundance biomass estimates. All other localities had less than 2% difference in proportions between MSA and target abundance female biomass estimates (Table S6).

## Discussion

We have designed a new GT-seq panel of 180 SNPs for genetic stock identification within the large Teno River Atlantic salmon fishery. The full panel performs equivalently to the previous microsatellite panel, with advantages in terms of cost and speed of genotyping, transferability among laboratories and flexibility for adding additional markers of interest. For example, in 2024 with an established pipeline and using a desktop short-read sequencer (Illumina MiSeq), generating the 180 SNP genotypes from 1,000 DNA extracts is estimated to require five days of staff time and €3,000 of laboratory consumables. It will also be more robust to anticipated future changes in genotyping technologies – in particular, greatly reduced availability of platforms and reagents currently used for microsatellite typing. Smaller sub-panels of the same SNPs also show good performance indicating that the full panel will be robust to a proportion of missing genotypes, a common outcome in GT-seq due to batch effects causing insufficient read depth over some targeted sites.

Our re-investigation of population structure within the Teno River, for which we re-genotyped archived samples for a large set of genome-wide SNPs, recapitulated the patterns previously found using microsatellites. Examination of genome-wide variation can reveal population genetic substructure that was not previously identified in studies using a smaller number of markers (reviewed in Sunde et al. 2020). However, this was not the case for our study - the existing 33 microsatellites appear to have been sufficient to resolve the population genetic substructure within the Teno River Atlantic salmon population. We caution against a general assumption that examining more variation across the genome will resolve increasingly more ecological- and management-relevant substructure within a target species.

Vähä *et al*. (2008), genotyping archived scale samples, demonstrated that this broad-scale population substructure in Teno River Atlantic salmon had remained stable over a minimum of four generations. Our results also support this, albeit on a smaller temporal scale: samples collected from the same site 3-6 years apart almost invariably clustered together in the Admixture analysis and PCA. Such temporally stable substructure is expected to arise and be perpetuated by the natal homing of salmon to their birth river. This in turn can enable the establishment of local adapted genotypes, which further reduces gene flow among populations by reducing the fitness of migrants. There is both genetic and demographic evidence for local adaptation within the Teno River salmon stock (Mobley *et al*., 2019; Pritchard *et al*., 2018), further emphasizing the importance of population-specific managing the fishery to avoid over-exploitation of certain subpopulations.

Despite this broad-scale stability of population structure, Vähä *et al*. (2008) did document change in allele frequencies across timepoints particularly in smaller tributary populations. We recommend additional juvenile collections to augment the baseline presented here, both to increase the sample size for poorly represented reporting units and to ensure that it accurately represents the contemporary genetic make-up of the Teno River salmon stock. A long-term monitoring programme should include plans to systematically update the baseline at pre-planned intervals and re-test the performance of the 180 SNP panel against the modified baseline.

Based on our examination of population structure, we chose to use 22 reporting units for the SNP-based GSI, fewer than the maximum of 32 that were used by (Vähä *et al*., 2017). Specifically, we amalgamated Ylakongas with the Teno River mainstem reporting unit, Luovtejohka with Tana Bru, Báišjohka with Nilijoki, Upper Iešjohka with Lower Iešjohka, and Iskurasjoki, Goššjohka, Vuomajoki and Cášcemjohka with the Inarijoki mainstem. Additionally, due to sample loss our baseline did not include two reporting units, Bavttájohka – a tributary of Kárášjohka and Anarjohka – a headwater tributary of Inarijoki. Nevertheless, investigation of the performance of the 180 SNP panel and the 33 microsatellite panel with the baseline divided into either 22 or 30 reporting units showed very similar results. Both demonstrated better overall assignment accuracy when there were fewer reporting units, while the SNPs actually performed better than the microsatellites for Báišjohka, Niljohka, Lower Iešjohka, Iskurasjoki and Goššjohka. Due to DNA degradation, our SNP baseline omitted two reporting units used by Vähä *et al*.(2017): Kárášjohka tributary Bavttájohka and Inarijoki tributary Anarjohka. Given the known patterns of genetic structure, we expect fish originating from these locations to be misassigned to a Kárášjohka/Iesjohka and an Inarijoki reporting unit, respectively. Bavttájohka contributes 1.8% of the target female biomass for the Teno River, while the biomass of Anarjohka is included in the Inarijoki and not calculated separately (Falkegård et al. 2014); thus, while we recommend addition of these locations into the SNP baseline we do not anticipate that their current absence is substantially biasing our results.

Average accuracy rates for other Atlantic salmon GSI systems range from 87% (33 microsatellites, 182 rivers in the northernmost Norway and Finland and in the northwest of Russia; (Ozerov *et al*., 2017) and between the upper and lower reaches of a large Baltic salmon river system (Miettinen et al. 2024), to 53.1 % at the level of the individual river and 72.1% at the regional level (31 microsatellite markers, 14 Norwegian rivers (Harvey *et al*., 2019). Considering the spatial scale of the Teno River system, the resolution using this GSI system is very high. Further increases in accuracy would be possible by merging rivers into broader, regional, reporting units, however, for management purposes tributary-based reporting units are more informative.

We caution that results from the microsatellites and SNP panel are not entirely comparable. In addition to differences in baseline sample sizes, the panel of GSI SNPs were selected and tested using the same individuals, an ascertainment bias that is expected to somewhat over-estimate their true assignment efficacy. We used this approach instead of dividing the baseline into training and test datasets in order to more accurately capture the true population allele frequency where we had low numbers of samples. Nevertheless, the proportion of the Teno River 2018 salmon catch assigned to the 22 baseline reporting units in the Mixed Stock Analysis closely scaled with the target female spawning biomass of those reporting units, suggesting that the SNP panel is indeed performing well: only three comparisons deviated more than 2% from the expected. Based on the comparison in this study alone, the outtake of fish in 2018 reflected quite accurately the potential size in female biomass of the populations (Falkegård *et al*., 2014) and the stock assessment status (Anon. 2024).

The Teno River 180 SNP panel, being easily modified with the addition of new SNPs, also has the potential to form the foundation of a more broadly applicable panel that could replace the current microsatellite panel for Genetic Stock Identification of Atlantic salmon throughout western Europe (Gilbey et al. 2021).

## Acknowledgements

This project was funded by LUKE, NINA and the Academy of Finland. We thank Teno/Tana/Deatnu River fishers for providing scale samples and Reeta Partanen for laboratory assistance. We also thank the numerous field workers collecting the Teno River baseline samples.

## Contributions

Project conceptualization: CRP, VLP, HJ, JE, MF, PO

Project management/Sample coordination: - PO, MF, AF, JE, CRP, VLP

Methods development: VLP

Laboratory analysis: VLP, HJ

Bioinformatics: VLP

Data analysis: HJ, VLP

Manuscript drafting: HJ, CRP, VLP

Visualization: HJ, VLP

Manuscript finalization: all authors.

## References

Ackerman, M. W., Habicht, C., & Seeb, L. W. (2011). Single-Nucleotide Polymorphisms (SNPs) under Diversifying Selection Provide Increased Accuracy and Precision in Mixed-Stock Analyses of Sockeye Salmon from the Copper River, Alaska. Transactions of the American Fisheries Society, 140, 865–881.

Alexander, D. H., Novembre, J., & Lange, K. (2009). Fast model-based estimation of ancestry in unrelated individuals. Genome Research, 19, 1655–1664.

Anderson, E. C. (2010). Assessing the power of informative subsets of loci for population assignment: standard methods are upwardly biased. Molecular Ecology Resources, 10, 701–710.

Anderson, E. C., Waples, R. S., & Kalinowski, S. T. (2008). An improved method for predicting the accuracy of genetic stock identification. Canadian Journal of Fisheries and Aquatic Sciences, 65, 1475–1486.

Anon. 2024. Status of the Tana/Teno River salmon populations in 2023. Report from the Tana/Teno Monitoring and Research Group nr 1/2024. Available at https://www.luke.fi/en/tana_research_group

Archer, F. I., Adams, P. E., & Schneiders, B. B. (2017). STRATAG: An R package for manipulating, summarizing and analysing population genetic data. Molecular Ecology Resources, 17, 5–11.

Aykanat, T., Lindqvist, M., Pritchard, V. L., Primmer, C. R., Lindqvist, M., & Primmer, C. R. (2016). From population genomics to conservation and management: a workflow for targeted analysis of markers identified using genome-wide approaches in Atlantic salmon Salmo salar. Journal of Fish Biology, 89, 2658–2679.

Barson, N. J., Aykanat, T., Hindar, K., Baranski, M., Bolstad, G. H., Fiske, P., … Primmer, C. R. (2015). Sex-dependent dominance at a single locus maintains variation in age at maturity in salmon. Nature, 528, 405–408.

Begg, G. A., Friedland, K. D., & Pearce, J. B. (1999). Stock identification and its role in stock assessment and fisheries management: an overview. Fisheries Research, 43, 1–8.

Campbell, N. R., Harmon, S. A., & Narum, S. R. (2015). Genotyping-in-Thousands by sequencing (GT-seq): A cost effective SNP genotyping method based on custom amplicon sequencing. Molecular Ecology Resources, 15, 855–867.

Chang, C. C., Chow, C. C., Tellier, L. C. A. M., Vattikuti, S., Purcell, S. M., & Lee, J. J. (2015). Second-generation PLINK: Rising to the challenge of larger and richer datasets. GigaScience, 4, 1–16.

Crozier, W. W., Schön, P.-J., Chaput, G., Potter, E. C. E., Maoileidigh, N. O., & MacLean, J. C. (2004). Managing Atlantic salmon (Salmo salar L.) in the mixed stock environment: challenges and considerations. ICES Journal of Marine Science, 61, 1344–1358.

Erkinaro, J., Niemelä, E., Vähä, J.-P., Primmer, C.R., Brørs, S. & Hassinen, E. 2010. Distribution and biological characteristics of escaped farmed salmon in a major subarctic salmon river. Implication for monitoring. Canadian Journal of Fisheries and Aquatic Sciences 67: 130–142.

Erkinaro, J., Czorlich, Y., Orell, P., Kuusela, J., Falkegård, M., Länsman, M., … Niemelä, E. (2019). Life history variation across four decades in a diverse population complex of Atlantic salmon in a large subarctic river. Canadian Journal of Fisheries and Aquatic Sciences, 76, 42–55.

Falkegård, M., Foldvik, A., Fiske, P., Erkinaro, J., Orell, P., Niemelä, E., … Hindar, K. (2014). Revised first-generation spawning targets for the Tana/Teno river system. 1–68 pp. Vol. 1087.

Fischer, M. C., Rellstab, C., Leuzinger, M., Roumet, M., Gugerli, F., Shimizu, K. K., … Widmer, A. (2017). Estimating genomic diversity and population differentiation - an empirical comparison of microsatellite and SNP variation in Arabidopsis halleri. BMC Genomics, 18, 1–15.

Frichot, E., & Francois, O. (2015). LEA: An R package for landscape and ecological association studies. Methods in Ecology and Evolution, 6, 925–929.

Gilbey, J., Utne, K. R., Wennevik, V., Beck, A. C., Kausrud, K., Hindar, K., De Leaniz, C. G., Cherbonnel, C., Coughlan, J., Cross, T. F., Dillane, E., Ensing, D., Jacobsen, J. A., Jensen, A. J., Karlsson, S., Maoiléidigh, N. Ó., Mork, A., Nielsen, E. E., Nøttestad, L., … Mcginnity, P. (2021). The early marine distribution of Atlantic salmon in the North-east Atlantic: A genetically informed stock-specific synthesis. Fish and Fisheries, 22, 1274–1306.

Harvey, A. C., Quintela, M., Glover, K. A., Karlsen, Ø., Nilsen, R., Skaala, Ø., … Wennevik, V. (2019). Inferring Atlantic salmon post-smolt migration patterns using genetic assignment. Royal Society Open Science, 6, 190426.

Kamvar, Z. N., Tabima, J. F., & Grünwald, N. J. (2014). Poppr: an R package for genetic analysis of populations with clonal, partially clonal, and/or sexual reproduction. PeerJ, 2, e281.

Koljonen, M.-L., & McKinnell, S. (1996). Assessing seasonal changes in stock composition of Atlantic salmon catches in the Baltic Sea with genetic stock identification. 998–1018 pp. Vol. 49.

Koljonen, M., Pella, J. J., & Masuda, M. (2005). Classical individual assignments versus mixture modeling to estimate stock proportions in Atlantic salmon (Salmo salar) catches from DNA microsatellite data. 2158, 2143–2158.

Larson, W. A., Seeb, J. E., Pascal, C. E., Templin, W. D., & Seeb, L. W. (2014). Single-nucleotide polymorphisms (snps) identified through genotyping-by-sequencing improve genetic stock identification of chinook salmon (oncorhynchus tshawytscha) from western alaska. Canadian Journal of Fisheries and Aquatic Sciences, 71, 698–708.

Li, H., & Barrett, J. (2011). A statistical framework for SNP calling, mutation discovery, association mapping and population genetical parameter estimation from sequencing data. 27, 2987–2993.

Li, H., & Durbin, R. (2010). Fast and accurate long-read alignment with Burrows–Wheeler transform. Bioinformatics, 26, 589–595.

Martin, M. (2011). Cutadapt removes adapter sequences from high-throughput sequencing reads. EMBnet Journal, 17, 10–12.

May, S. A., McKinney, G. J., Hilborn, R., Hauser, L., & Naish, K. A. (2020). Power of a dual-use SNP panel for pedigree reconstruction and population assignment. Ecology and Evolution, 10, 9522–9531.

Miettinen, A., Romakkaniemi, A., Dannewitz, J., Pakarinen, T., Palm, S., Persson, L., Östergren, J., Primmer, C. R., & Pritchard, V. L. (2024). Temporal allele frequency changes in large-effect loci reveal potential fishing impacts on salmon life-vhistory diversity. Evolutionary Applications, 17, e13690.

Mobley, K. B., Granroth-Wilding, H., Ellmen, M., Vähä, J.-P., Aykanat, T., Johnston, S. E., … Primmer, C. R. (2019). Home ground advantage: Local Atlantic salmon have higher reproductive fitness than dispersers in the wild. Science Advances, 5, eaav1112.

Moran, B. M., & Anderson, E. C. (2019). Bayesian inference from the conditional genetic stock identification model. Canadian Journal of Fisheries and Aquatic Sciences, 76, 551–560.

Ozerov, M., Vähä, J.-P., Wennevik, V., Niemelä, E., Svenning, M.-A., Prusov, S., … Christiansen, B. (2017). Comprehensive microsatellite baseline for genetic stock identification of Atlantic salmon (Salmo salar L.) in northernmost Europe. ICES Journal of Marine Science, 74, 2159–2169.

Palsbøll, P. J., Bérubé, M., & Allendorf, F. W. (2007). Identification of management units using population genetic data. Trends in Ecology and Evolution, 22, 11–16.

Paradis, E., & Schliep, K. (2019). Phylogenetics ape 5.0: an environment for modern phylogenetics and evolutionary analyses in R. Bioinformatics, 35, 526–528.

Parrish, D. L., Behnke, R. J., Gephard, S. R., McCormick, S. D., & Reeves, G. H. (1998). Why aren’t there more Atlantic salmon (Salmo salar)? Canadian Journal of Fisheries and Aquatic Sciences, 55, 281–287.

Prevosti, A., Ocana, J., & Alonso, G. (1975). Distances between Populations of Drosophila subobscura, Based on Chromosome Arrangement Frequencies. Theoretical and Applied Geentics, 45, 231–241.

Pritchard, V. L., Mäkinen, H., Vähä, J. P., Erkinaro, J., Orell, P., & Primmer, C. R. (2018). Genomic signatures of fine-scale local selection in Atlantic salmon suggest involvement of sexual maturation, energy homeostasis and immune defence-related genes. Molecular Ecology, 27, 2560–2575.

R Core Team. (2014). R: A language and environment for statistical computing. R Foundation for Statistical Computing, Vienna, Austria. URL http://www.R-project.org/.

Storer, C. G., Pascal, C. E., Roberts, S. B., Templin, W. D., Seeb, L. W., & Seeb, J. E. (2012). Rank and Order: Evaluating the Performance of SNPs for Individual Assignment in a Non-Model Organism. PLoS ONE, 7, 49018.

Sunde J, Yildirim Y, Tibblin P, Forsman A. Comparing the Performance of Microsatellites and RADseq in Population Genetic Studies: Analysis of Data for Pike (Esox lucius) and a Synthesis of Previous Studies. Front Genet. 2020 Mar 13;11:218. doi: 10.3389/fgene.2020.00218. PMID: 32231687; PMCID: PMC7082332.

Untergasser, A., Cutcutache, I., Koressaar, T., Ye, J., Faircloth, B. C., Remm, M., & Rozen, S. G. (2012). Primer3-new capabilities and interfaces. Nucleic Acids Research, 40, e115.

Vähä, J.-P., Erkinaro, J., Niemelä, E., Primmer, C. R., Saloniemi, I., Johansen, M., … Brørs, S. (2011). Temporally stable population-specific differences in run timing of one-sea-winter Atlantic salmon returning to a large river system. Evolutionary Applications, 4.

Vähä, J.-P., Erkinaro, J., Falkegård, M., Orell, P., & Niemelä, E. (2017). Genetic stock identification of Atlantic salmon and its evaluation in a large population complex. Canadian Journal of Fisheries and Aquatic Science, 74, 327–338.

Vähä, J.-P., Erkinaro, J., Niemelä, E., & Primmer, C. R. (2007). Life-history and habitat features influence the within-river genetic structure of Atlantic salmon. Molecular Ecology, 16, 2638–2654.

Vähä, J.-P., Erkinaro, J., Niemelä, E., & Primmer, C. R. (2008). Temporally stable genetic structure and low migration in an Atlantic salmon population complex: implications for conservation and management. Evolutionary Applications, 1, 137–154.

Winans, G. A., Paquin, M. M., Van Doornik, D. M., Baker, B. M., Thornton, P., Rawding, D., … Kalinowski, S. (2004). Genetic Stock Identification of Steelhead in the Columbia River Basin: An Evaluation of Different Molecular Markers; Genetic Stock Identification of Steelhead in the Columbia River Basin: An Evaluation of Different Molecular Markers. North American Journal of Fisheries Management 24, 672–685.

Yano, A., Nicol, B., Jouanno, E., Quillet, E., Fostier, A., Guyomard, R., & Guiguen, Y. (2013). The sexually dimorphic on the Y-chromosome gene (sdY) is a conserved male-specific Y-chromosome sequence in many salmonids. Evolutionary Applications, 6, 486–496.

